# Cont-ID: Detection of samples cross-contamination in viral metagenomic data

**DOI:** 10.1101/2023.01.23.525161

**Authors:** Johan Rollin, Wei Rong, Sébastien Massart

**Affiliations:** University of Liège, Gembloux Agro-Bio Tech, Plant Pathology Laboratory, 5030, Gembloux, Belgium; DNAVision, 6041, Gosselies, Belgium

**Keywords:** Bioinformatic, virus, detection, sequencing, contamination, metagenomic

## Abstract

**Background:** High Throughput sequencing (HTS) technologies completed by the bioinformatic analysis of the generated data are becoming an important detection technique for virus diagnostics. They have the potential to replace or complement the current PCR-based methods thanks to their improved inclusivity and analytical sensitivity, as well as their overall good repeatability and reproducibility. Cross-contamination is a well-known phenomenon in molecular diagnostics and corresponds to the exchange of genetic material between samples. Cross-contamination management was a key drawback during the development of PCR-based detection and is now adequately monitored in routine diagnostics. HTS technologies are facing similar difficulties due to their very high analytical sensitivity. As a single viral read could be detected in millions of sequencing reads, it is mandatory to fix a detection threshold that will be influenced by cross-contamination. Cross-contamination monitoring should therefore be a priority when detecting viruses by HTS technologies.

**Results:** We present Cont-ID, a bioinformatic tool designed to check for cross-contamination by analysing the relative abundance of virus sequencing reads identified in sequence metagenomic datasets and their duplication between samples. It can be applied when the samples in a sequencing batch have been processed in parallel in the laboratory and with at least one external alien control. Using 273 real datasets, including 68 virus species from different hosts (fruit tree, plant, human) and several library preparation protocols (Ribodepleted total RNA, small RNA and double stranded RNA), we demonstrated that Cont-ID classifies with high accuracy (91%) viral species detection into (true) infection or (cross) contamination. This classification raises confidence in the detection and facilitates the downstream interpretation and confirmation of the results by prioritising the virus detections that should be confirmed.

**Conclusions:** Cross-contamination between samples when detecting viruses using HTS can be monitored and highlighted by Cont-ID (provided an alien control is present). Cont-ID is based on a flexible methodology relying on the output of bioinformatics analyses of the sequencing reads and considering the contamination pattern specific to each batch of samples. The Cont-ID method is adaptable so that each laboratory can optimise it before its validation and routine use.

## Background

The advent of high-throughput sequencing (HTS) technologies coupled with the development of powerful bioinformatics approaches has improved our ability to detect viruses in a non-targeted way from any sample collected from diverse sources. Noteworthy, detecting viruses by HTS technologies relies on many steps in the laboratory: sampling, transport and storage, nucleic acid extraction, library preparation, and sequencing (1). Compared to other molecular tests like (RT)-PCR, these steps are much more numerous and complex (2).

The analytical sensitivity, e.g. the ability to detect viral species at very low concentration in a sample, has been demonstrated to be similar to or even better than RT-PCR for animals (3) or plant viruses(4,5)). In addition, the inclusivity of HTS technologies, e.g. the ability to detect all isolates from a species and all species whose nucleic acids are present in enough quantity in a nucleic acid extract, is particularly high compared to any other detection test (2,6). Consequently, the use of HTS technologies is currently expanding at a rapid pace in research and is also progressively used for the diagnostic of viruses threatening humans (7), including SARS-Cov-2 (8), livestock (9) or plant health (10).

The broader application of HTS technologies for virus detection, with the simultaneous analysis of tens to hundreds of samples, is raising a significant challenge that needs to be addressed: the management of cross-contamination between samples. Scientists and diagnosticians already faced such challenges decades ago during the development of PCR-based techniques for detecting plants (11,12) or animal viruses (13,14), and this phenomenon might worsen with the use of HTS for virus detection (2). The higher complexity of laboratory operations, the intrinsically very high inclusivity, and the very low limit of detection (few viral reads are enough to detect the virus) of HTS make cross-contamination a more pressing issue. This is a frequently observed but, until recently, rarely reported observation in many, if not all, laboratories that have tested these technologies for virus detection. In many cases, these problems are frequently limited to a low number of reads and are of little consequence. Still, the specifics of the diagnostics field, with the need to detect viruses that can be at very low titre in the sample, clearly give more impact to such potential contamination problems (2). The occurrence of contamination is, therefore, a key element to consider when interpreting the viruses detected in HTS datasets.

The consequences of erroneous detection due to cross-contamination between samples can be catastrophic, as described for tuberculosis prior to HTS (15) but also using HTS for human and plant viruses (16,17).

So, even if the cross-contamination issue of HTS is long known and discussed in the scientific community, proper methodologies and dedicated algorithms are still missing to address it. Until now, the burden of detection confirmation relied on the virologist’s expertise and the use of laboratory tests to independently confirm the presence of the virus in the sample, which is a fastidious, costly, and time-consuming task. To minimise the confirmation burden, arbitrary thresholds (like 5 or 10 reads) (4,18) have been proposed to consider a detection valid. Still, these thresholds are subjectively fixed based on the sequencing/detection tools or the scientist’s experience. In addition, it has been shown recently that the cross-sample contamination burden can be very variable between sequencing batches and that an adaptative threshold is required(5). Therefore, the need for formal bioinformatic pipelines for HTS-based data that consider the possible cross-contamination is growing (19).

To handle cross-contamination, several laboratory protocol improvements have been implemented over time: laboratory or reagent decontamination, alternate dual indexes, inter-run washing (20,21) or, more recently, the use of alien control. An alien control is defined as “a matrix infected by a target (called alien target) which belongs to the same group as the target organism to be tested in the samples, but that cannot be present in the samples of interest.” (22). It is processed as external control alongside the sample to be analysed. It is preferably the same type of matrix as the analysed samples: plant tissue, water … Ideally, the alien target, in our case a virus, should be at a high concentration in the alien sample as it allows a better analysis of cross-contamination between samples. Indeed, the probability of detecting any virus at a low level due to cross-contamination rises if this virus is very abundant in at least one of the processed samples. A high abundance of the alien virus will therefore allow better monitoring of contaminations, including for other viruses highly abundant in at least one tested sample. The presence of sequencing reads from the alien virus in any tested sample can be considered the consequence of contamination from the alien control to this sample. Such information can be used to monitor the cross-contamination level between samples within the sequencing batch.

Many generalist bioinformatic tools, such as Kraken (23) or BLAST (24) can detect the presence of viruses in HTS datasets with very high analytical sensitivity, as the detection is possible from a single viral read or contig. Some of them, like VirHunter (25), VirAnnot (26) or VirusDetect (27), have been specifically developed for that purpose. Nevertheless, they have not been designed to detect cross-contamination in the input datasets. Instead, they will detect a virus, whatever its origin: virus infection in the biological sample or contamination from another sample. The risk of contamination is particularly acute for viruses in very high abundance in one of the samples sequenced as a few contaminating reads can be detected by the bioinformatic tools in other samples prepared in parallel. The situation’s impact is growing, especially in the diagnostic field (2), as false positive results due to contamination can lead to inaccurate data interpretation, which can cause tremendous health and trade issues.

According to EDAM ontology (28), tools that address cross-contamination issues should be labelled as “Sequence contamination filtering”. We were looking for tools using EDAM terms and the usual ones (virus reads contamination, cross-contamination, …). Some tools address similar issues like contamination on bacterial isolates (ConFindr- (29) or bacterial metagenome (GUNC - (30). They both use methods relying on operons organisation of genes that are not applicable for viruses. Croco (31) uses an approach mainly based on bacterial quantitative data. Finally, DecontaMiner (32) can be applied to metagenome data, including viruses but is based on a combination of detection methods (mainly mapping and Blast) that try to assign the dark matter (reads from unknown origin) more than formally detecting the cross-contamination material. To our knowledge, there is no tool specifically addressing cross-contamination during virus detection in metagenome datasets. It means that some risks of false positive results remain unmonitored for virologists, and the burden of confirmation of detection in case of false positive is still not addressed.

To solve this issue, we present Cont-ID, a method designed to check sample cross-contamination for viruses previously identified in metagenomic datasets. It relies on a simple requirement: every sample in a sequencing batch should have been processed at the same time and followed the same steps in the laboratory with at least one alien control as external control. Cont-ID uses a voting system to classify every species prediction on each sample of the sequencing batch into (true) infection or (cross) contamination. This tool will help the virologist to distinguish virus presence and virus cross-contamination in HTS data improving the reliability of viral detection and the efficiency of downstream confirmation and characterisation analyses. It can also help to improve feedback on upstream steps that might be linked to cross-contamination events. Cont-ID is an open-source python (v3) based script method freely available here: https://github.com/johrollin/viral_contamination.

## Methods

### Implementation

In viral metagenomics, detecting multiple viral species in the same sample is frequent, and a virus species can be seen with different confidence levels in several samples of the same sequencing batch. Therefore, Cont-ID aims to determine whether a given detected virus in a sample is likely to be a contaminant or not by comparing it to the results from the other samples of the same sequencing batch, e.g. samples processed in parallel and following the same laboratory steps.

Cont-ID does not require any development or maintenance of database as it only relies on data generated by usual bioinformatics tools for HTS dataset analyses and, most specifically, two elements: (i) the normalised abundance estimation (number of reads assigned to each detected virus species on each sample) and (ii) the number of identical reads among pairs of samples (deduplication ratio). These input metrics are easy to obtain as the abundance estimation can be calculated by using any mapping tool like BWA (33) or a read classifier like Kraken/Bracken (23,34), and the number of identical reads from a virus between two samples can be obtained by running any deduplication tool like BBduck (35). A tabulated file containing these numbers associated with the detected virus name and the unique ID for each batch sample is used as input for Cont-ID, as shown in **Figure 2**. Each virus predicted on each batch sample is considered a distinct element and corresponds to a line in the generated table. A separate table is generated for the alien virus.

Computing the two elements mentioned above into three different metrics for the alien virus and each detected virus, Cont-ID can predict through three rules if a given viral species detection is likely a cross-contaminant or not in the sequencing batch, as described in **Figure 1**.

**Figure 1:**
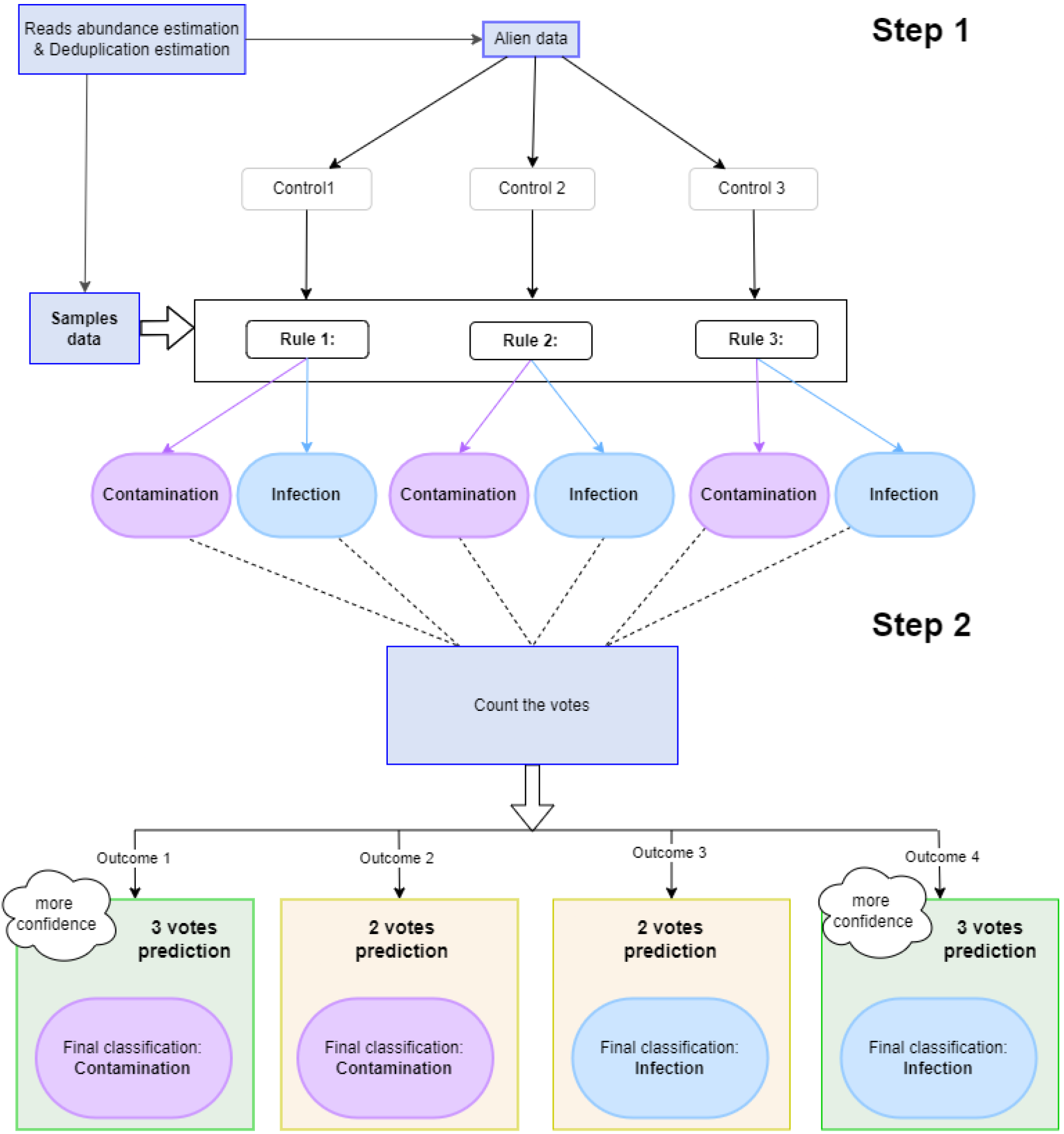
Cross-contamination prediction with Cont-ID. There are two input files, one for the alien data and one for the samples data. Alien file is used to calculate control thresholds which are then used along with the sample data to apply rules to a voting system (step 1). The votes are then counted to decide for each virus on each sample (element) either if it is a (cross)contamination or an infection (step 2).

**Figure 2:**
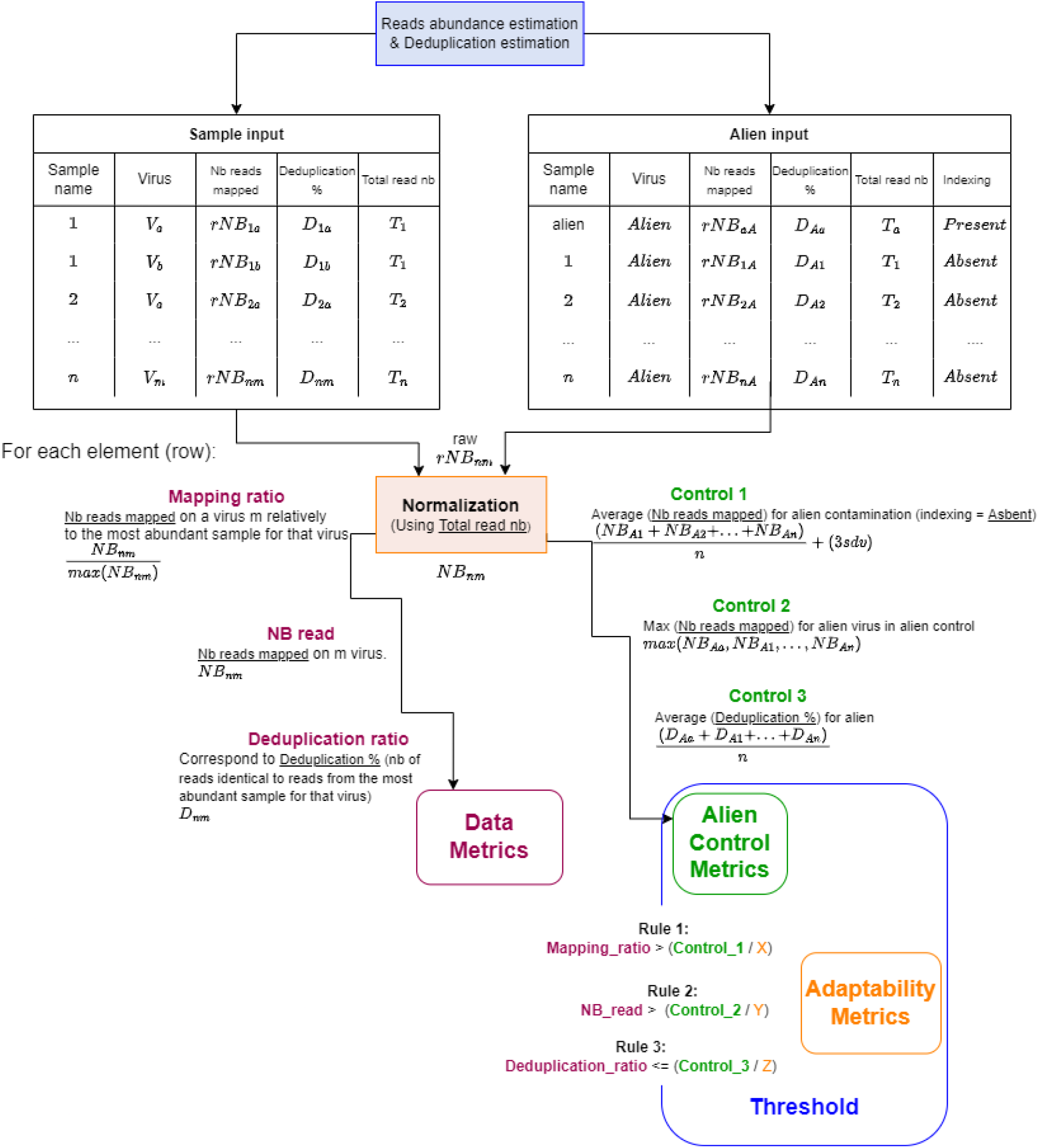
Cont-ID rules explanation. There are two input files, one for the alien data and one for the samples data. Alien file is used to calculate each alien control metric after normalisation. The sample file is used to calculate each data metric after normalisation. Each alien control metric is associated with a user (manually) designed adaptability metric (X, Y or Z) to compose each rule’s threshold. Finally, each Data Metric is compared to the corresponding threshold in order to obtain the three rules used in Cont-ID.

The three rules classify as contamination or infection each element according to the pattern of reads number observed among the samples and the alien control for the alien virus and the considered viral species. Rules one and two both use the (normalised) reads abundance estimation, while rule three uses the assessment of unique (identical) reads. Rules are calculated after normalising the number of reads per sample and are described more precisely in **Figure 2**.

The first rule uses the mapping ratio of each virus in each sample (corresponding to an element): the number of reads of each element is divided by the maximum number of reads of the corresponding virus in one of the samples. This first rule compares this mapping ratio for the element with the Control 1 metrics calculated for the alien virus and corresponds to the average number of reads mapped on the alien virus in the samples for which the alien is a cross-contaminant, with three times the standard deviation of this average.

The second rule relies on the number of reads of the element in the sample. The rule compares it with Control 2, corresponding to the number of alien reads identified in the alien control. The third rule is based on each element’s deduplication ratio, which is compared with Control 3. The average deduplication ratio of the alien virus reads between each tested sample and the alien control.

We aimed to find the most reliable formula for threshold calculation on each rule while allowing a part of adaptability according to the biological system used. As the system variability can come from the laboratory using HTS, the host and type of sample (fruit tree, herbaceous plant, human, animal …), the type of virus (integrated or non) or the extraction protocol used (dsRNA, total RNA, small RNA…), each rule includes a third number (represented by X, Y or Z) that is called adaptability metrics (see **Figure 2**). The X will impact the first rule that considers the relative proportion of reads of a virus in this sample compared to the sample with the maximum read of this virus. This threshold is a refinement of the “alien threshold” described earlier(5). The default value proposed is 2. The Y divides the number of reads from the alien virus in the alien control for comparing it to the number of reads of each virus in each sample. In this publication, a default value of 1,000 has been fixed for Y, and it was in the range of the expected (cross) contamination ratio (number of reads in the truly infected sample versus the number of reads in contamination one). The Z metric impacts Control 3 and the evaluation of the proportion of identical reads between different samples. The proportion of identical reads can be influenced by different factors (mutation rate, respective genome length …). The role of Z is to consider those different factors. A default value of 1.5 is proposed.

Default values of the three adaptability metrics have been provided in this publication after their optimisation on the banana datasets and their evaluation of other datasets. Nevertheless, users can independently modify them during the evaluation or validation of Cont-ID applied to their datasets. A careful evaluation of the adaptability metrics by the user is recommended to evaluate their impact on the diagnostic performance of the test. In addition, several sets of adaptability metrics can be run in parallel for further improvements in diagnostics performance. The value given to the adaptability metrics and controls resulting is always recorded in an additional log file (see Supplementary File 1 [log_file]). This log file help to ensure traceability allowing the user to check the pertinence of the chosen numbers and to adapt them when needed. As each of the three rules has two possible decisions (contamination or infection), a majority vote will be obtained with two or three votes. The decision of each vote is available in the generated result to support the result interpretation and let the user decides on the confidence to give to each individual rule according to the biological system tested.

In addition, the proper quantitative comparison of sequencing reads datasets relies on normalising the number of reads per sample, for example, as always done for transcriptomic or microbiome studies. This assertion is also true for Cont-ID and corresponds to an adaptative parameter. To limit some bias due to the difference in sequencing depth between samples in the same batch, we also normalise by default to 5 000 000 reads in this publication. Still, it is manually changeable by the user.

Finally, Cont-ID also has another level of flexibility: the script is made to ease the change of rules in the code that can complete or replace the existing ones.

### Conditions of application for Cont-ID

Three main conditions are essential to run Cont-ID. First, an alien control should be used, alien control should contain a high concentration of the alien virus, so reads from that viral species are more prone to be detected in other samples when cross-contamination occurs from the alien control to the other samples. Similarly, if another virus is found in the alien control sample, that is also an indication of potential contamination (although not used so far by Cont-ID). The alien control is bioinformatically processed exactly as the samples of interest to generate the alien metrics for each sample (in a separate tabulated file). In the absence of external alien control, it is still possible to analyse sequencing batches if they include samples from different host species and some detected viruses, preferably at high abundance, are known to infect only some of the host species. In such a case, the alien file should be filled with the selected virus as if it was an alien (with the status of alien present/absent in the file). Nevertheless, the threshold set-up and the results will be less accurate and include fewer samples (the samples corresponding to species that can be hosts for the virus could not be considered). In addition, a high degree of confidence is needed regarding the actual infection of the sample selected as alien control by the virus selected as an alien virus. Cont-ID always requires at least one (cross) contamination in the alien file to be reported; otherwise, the threshold calculation will fail; in that case, the tool will state it.

The second application condition is related to the processing of the samples and the alien control. The alien control and all the other samples in a given batch should have been processed together in parallel for all the laboratory steps (RNA/DNA extraction, library preparation, sequencing) and bioinformatics (Reads cleaning, host removing …). This is a good diagnostic practice, but it is even more important here as the goal is to observe cross-contamination levels. The assumption is that the level seen with the alien represents what could have happened in samples of interest. Therefore, this assumption depends on processing all samples and control in parallel.

The third condition is that, once the user has fixed the adaptability metrics, the analysis should be carried out batch per batch. The calculation of sample and alien metrics is dynamically done for each batch as cross-contamination patterns can strongly vary between batches, as recently shown for banana samples (5).

### Sequencing reads datasets

The first datasets (batches A to D) were generated in our laboratory by total RNA sequencing protocol with ribodepletion applied to RNA extracted from banana plants (belonging to the *Musa* genus) (5). These data were generated to compare the test performance criteria of high throughput sequencing with classical virus testing protocols that include ImmunoCapture (IC)-(RT)-PCR and electron microscopy (36). The alien control corresponded to wheat plants infected by two species of barley yellow dwarf virus (BYDV-PAS and BYDV-PAV)(5). In total, four sequencing datasets (called A, B, C, and D) composed respectively of 27, 20, 27 and 25 samples were generated independently. A fifth batch generated during this validation experiment using diluted samples for evaluating the limit of detection (analytical sensitivity) was not included in our analysis according to the recent guidelines proposed for statistical analysis of validation datasets for plant pest detection (22). A total of 10 different viral species were infecting these samples, including banana mild mosaic virus (BanMMV), banana bract mosaic virus (BBrMV), banana bunchy top virus (BBTV), cucumber mosaic virus (CMV), and five species belonging to the banana streak virus (BSV) species complex. In addition, two other sequencing protocols were applied to some banana plants, small RNA sequencing (5) starting from the same RNA extract as total RNA sequencing (for 21 samples in a single batch) and double-stranded RNA (dsRNA) enrichment and sequencing protocol (37) applied from plant tissue of 13 samples in a single sequencing batch.

BSV is a species complex (genus: *Badnavirus*, family: *Caulimoviridae)* among which five species were included in our samples: banana streak CA virus (BSCAV), banana Goldfinger virus (BSGFV), banana streak IM virus (BSIMV), banana streak Mysore virus (BSMYV), and banana streak OL virus (BSOLV). Notably, some species of this complex have their genome fully or partially integrated into the plant genome as endogenous viral elements (EVE), most specifically in B genomes originating from *M. balbisiana.* These EVE can be transcribed in the plant, and for some BSV species, they can even trigger an infection with viral particles of BSV in the plant (38). It is well documented that BSGFV, BSIMV and BSOLV are constitutive of *Musa balbisiana* (B genome) but can be activated in some conditions (39). In addition, BSMyV is also integrated into the Musa B genome, although the ability to produce infectious viral particles is not yet demonstrated. This brings additional complexity as EVE can be transcribed without the presence of a viral particle. It has been recently demonstrated that the detection of BSV transcripts by HTS tests must be confirmed by an independent test such as immunocapture (IC)-PCR for confirming the presence of viral particles (5).

The other datasets used in this work came from publicly available datasets listed in **Table 1** and were already included in peer-review publications. They were selected because they fit two criteria: (i) having all virus presence checked in all the samples and (ii) having a virus species that could act like an alien control for the input file. First, another set generated to detect viruses from diverse plant samples by high throughput sequencing of total RNA extraction was kindly provided by Queensland University of Technology (17), corresponding to a total of 19 plant viruses and viroid in 5 samples. In addition, the datasets generated from human samples came from published data from 3 different sources, with a total of 129 samples containing 39 viral species (40–42). These three human datasets allowed us to test Cont-ID with a large diversity of viruses, with different extraction and sequencing methods listed in Supplementary File 2.

**Table 1:**
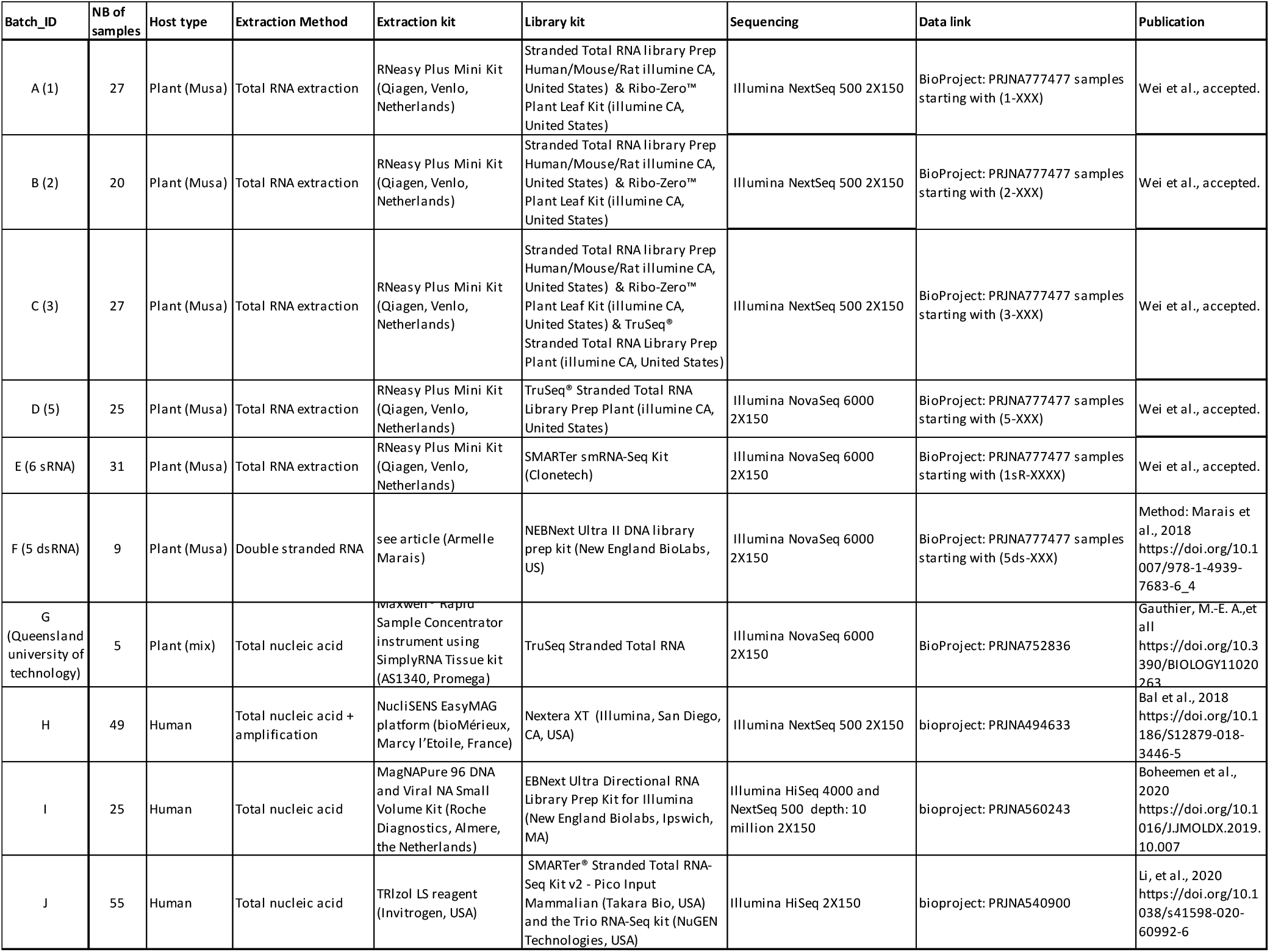
list of datasets used on Cont-ID

In total, ten sequencing batches, including 273 samples and the presence of 68 viral species, were used to test the potential impact of a different host, extraction, and sequencing method on Cont-ID performances. All the data generated are available with the link and procedure applied to obtain them described in **Table 1**; the indexing status of each virus in each sample is also available in Supplementary File 2.

### Bioinformatic analyses

#### Quality control and mapping of sequencing reads

For all datasets, read quality control (quality trimming, reads deduplication) was performed using a standard procedure described elsewhere (5). The cleaned reads were then mapped to a custom-built database (DB) containing all complete genome sequences from previously detected viruses in the datasets. For banana samples, all the complete genome sequences of the viruses were downloaded from NCBI nt database on (12/12/2020) to serve as mapping DB. While the BYDV reference (KU170668 – for the alien control) was selected as it was the closest sequence from our isolate. More information on the composition of each mapping DB is available elsewhere (5).

The reads were mapped on the custom DB using Geneious mapper (Prime 2020.0.5, Biomatters). First, the profile parameters “Low sensitivity / Fastest” were selected (with 20% mismatch and a maximum of 3 nucleotides gap allowed). To improve the results by aligning reads to each other in addition to the reference sequence, the fine-tuning for mapping was set to “Iterate 2 times”. The “multiple best matches” option was set to “Randomly” (no multiple best matches between two different viruses were observed in any sample processed). In the coming result section, we will refer to these parameters as “relax”. A second mapping referred to later on as “strict” was carried out using the same parameters except for the mismatch allowance that was lower than 10%. Only the second mapping was carried out for small RNA (20% mismatch is too much for small RNA). Indeed, using mismatches up to 20% should allow better inclusivity of the analysis by mapping reads from isolates that can be genetically distant from the reference sequences, especially if few reference genomes are available in the literature. Mapping with a strict parameter was done to use small RNA and confirm this hypothesis. The tolerance of mismatches of 20% is also close to many ICTV demarcation criteria to distinguish two different species (although these criteria are often considered for only one or a few genes and might vary between families). Another test with more relaxed parameters would increase the risk of adding non-specific reads (e.i. not generated from the viral genomes) and was not considered.

#### Deduplication of identical reads between samples

To investigate cross-contamination between samples, additional deduplication of identical reads between samples was performed using dedupe V38.37 (from BBMap) embedded in Geneious (Prime 2020.0.5, Biomatters) and with the parameters kmer seed length, maximum edit, and maximum substitutions set as “31”, “0”, and “0”, respectively. For each virus and sample, the mapped reads from each tested sample and the sample with the highest number of mapped reads in the batch were grouped into a single pool (using “Group sequences into a list” in Geneious) and deduplicated. The deduplication percentage equalled the number of reads removed as duplicates divided by the lower number of reads between the two tested samples. The deduplication percentage was not calculated on samples if less than 5 reads were mapped to target viruses. For those samples, the rule (number three) automatically votes contamination. While for the samples with the highest number of reads for a given virus, the deduplication ratio was set as reference (i.e. “RF”), and the vote for rule three is infection.

#### Confusion matrix and performance criteria calculation

We used a confusion matrix for each batch’s results to have standard metrics for comparing batches and samples. We compared the tool prediction for each element to the indexing status of the dataset assimilating infection as a positive result and contamination as a negative result, as explained in **Table 2**.

**Table 2:**
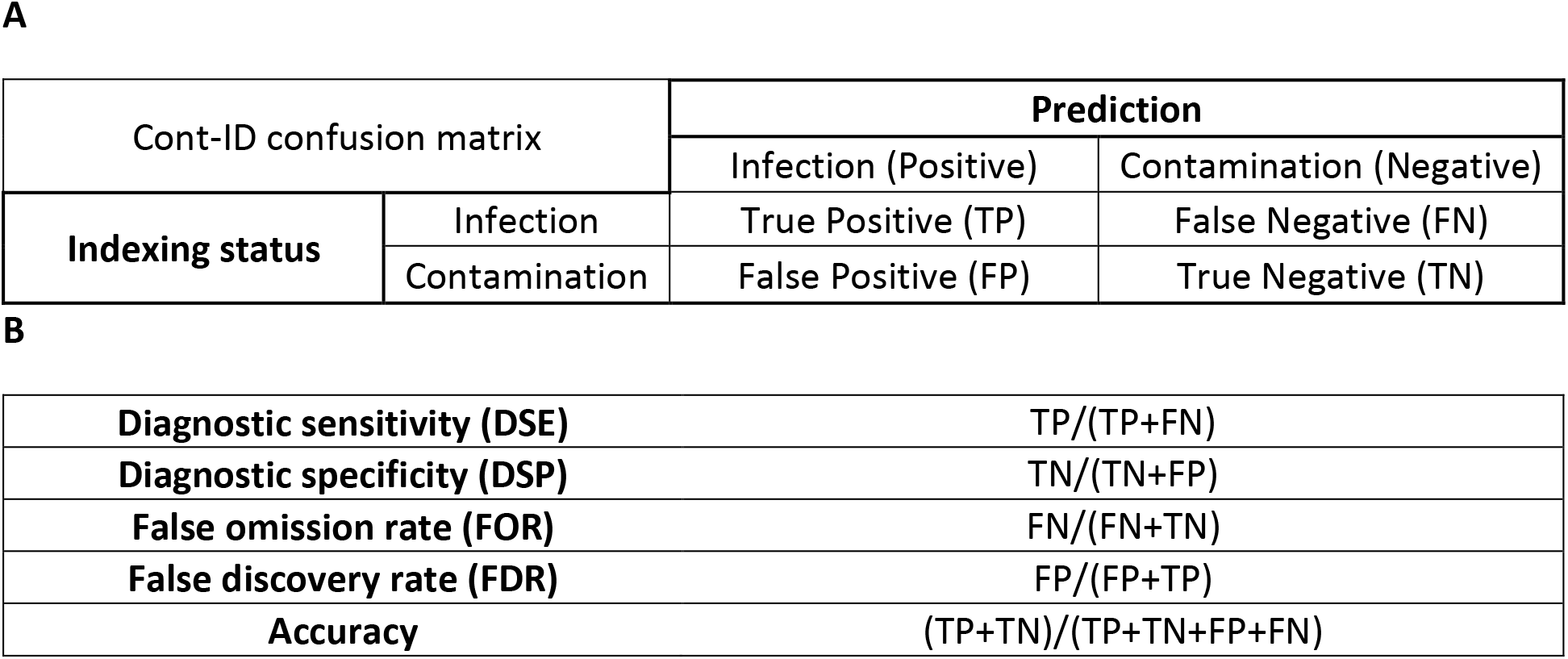
(A) Confusion matrix based on Cont-ID results. (B) The formula is used to calculate tool performance criteria.

| Cont-ID confusion matrix |  | Prediction |  |
| --- | --- | --- | --- |
|  |  | Infection (Positive) | Contamination (Negative) |
| Indexing status | Infection | True Positive (TP) | False Negative (FN) |
|  | Contamination | False Positive (FP) | True Negative (TN) |

|  |  |
| --- | --- |
| **Diagnostic sensitivity (DSE)** | $TP/(TP+FN)$ |
| **Diagnostic specificity (DSP)** | $TN/(TN+FP)$ |
| **False omission rate (FOR)** | $FN/(FN+TN)$ |
| **False discovery rate (FDR)** | $FP/(FP+TP)$ |
| **Accuracy** | $(TP+TN)/(TP+TN+FP+FN)$ |

Based on the confusion matrix, we have four possibilities after a prediction: False Positive (FP) when the tool wrongly predicted an infection, True Positive (TP) when the tool correctly predicted an infection, False Negative (FN) when the tool wrongly predicted contamination and True Negative (TN) when the tool correctly predicted contamination. In addition, we calculated several performance criteria commonly used in diagnostics to evaluate our tool. To calculate those performance criteria automatically, we used an automated script available on the same GitHub (https://github.com/johrollin/Cont_ID/tree/master/further_analysis).

## Results

We used Cont-ID on ten metagenomic datasets, including a total of 273 samples, as a proof of concept (see details in method). These datasets covered a broad range of use for Cont-ID as they were generated from plant or human samples according to three library preparation protocols (total RNA, small RNA and double stranded RNA).

### Set up adaptability metrics datasets on the banana datasets

When applying for the first time Cont-ID on banana datasets generated from reference samples with known viral status, the first objective was to determine the most appropriate values for the adaptability metrics (X, Y and Z), allowing to minimise both FP (over-prediction of infection) and FN (over-prediction of contamination). This was particularly complex as raising the value of an adaptability metric could lead to an over-prediction of either contamination or infection by the rule, while lowering it had the opposite effect.

During the set-up of the method, we looked for the most adapted set of values to balance our rule prediction on *Musa* datasets A, B and C. We tested several ranges of values aiming at limiting both wrong predictions (FP and FN). The optimised single set of values maintaining FP and FN low in the three datasets was not found. Indeed, variability was observed between batches, as any set limiting FP and FN in one or several batches was not optimal for the other batch(es).

Indeed, the uneven proportion and pattern of cross-contaminations observed in different sequencing batches made it very difficult to decide on a unique set of values. Instead, it seemed more efficient to apply two different sets of values (called “case 1” and “case 2” further on) that favoured the prediction of either true infection (TP - case 1) or true contamination (TN - case 2) from the same datasets. The combination of the prediction from both cases would give additional information for interpretation. We proposed values that gave the best performance criteria on our training datasets on bananas, and the purpose of the diagnostic test was to minimise the risk of false negatives (priority 1) while keeping the confirmation burden manageable (priority 2). Importantly, those sets of values can be manually adapted by the user to improve one or several performance criteria of the test, to better fit the purpose of the HTS tests carried out and its associated risks (risk of false positive or false negative) or to limit the “grey zone” of inconclusive results (see under).

Therefore, we propose to run Cont-ID with two sets of adaptability metrics every time to compare the results. Therefore, a high level of confidence is reached for the elements with identical predictions between both cases. The combination can also highlight elements for which the prediction changed; they correspond to the “grey zone” with metrics of abundance and/or duplication close to thresholds. In such cases, the automated prediction might not be accurate. At this stage, it is mandatory to carry on additional verification, such as checking the confidence (2 or 3 votes) for each prediction or comparing the threshold numbers (also provided in the result) with the sample metrics. Cont-ID provides the list of votes for each rule in each case to facilitate this additional verification. Then according to the additional information and the test’s purpose, the user can decide on the status (infection or contamination) or keep it inconclusive but decide to test the virus presence independently by another test. For the presentation of the result, the result is mentioned as “inconclusive” when both cases disagree.

### Evaluation of the method accuracy on banana samples

Based on the results obtained with the two sets of adaptability metrics, the tool predictions were compared with the biological status of each reference banana sample (batches A, B, C and D Supplementary File 2), allowing us to predict the cross-contamination on the four tested batches with an average accuracy of 90%, excluding 23% of elements classified as “inconclusive” (see table 3A).

**Table 3:**
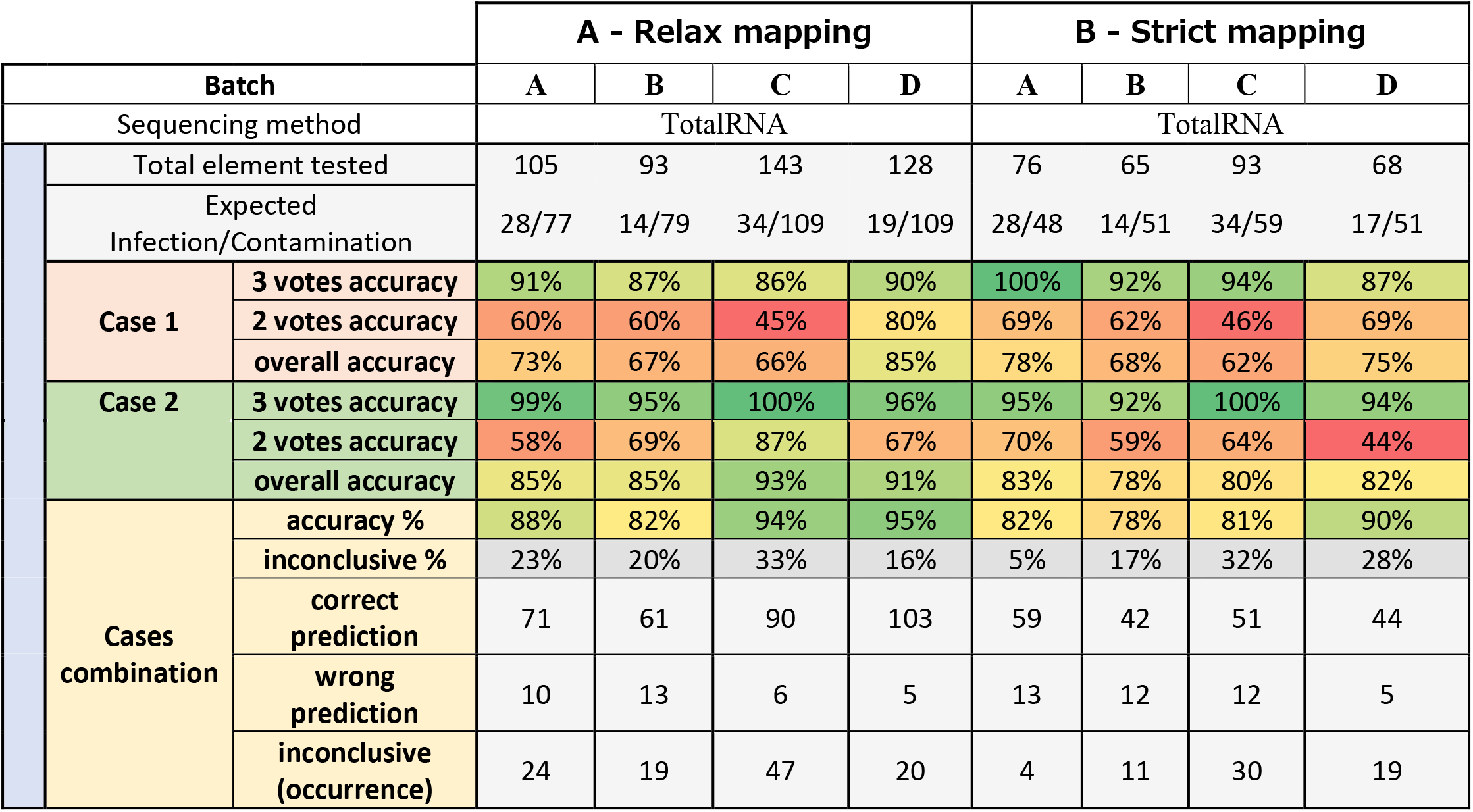
Percentage of the accuracy of case 1 and case 2 analysed alone or in combination on banana samples sequenced by ribodepleted totalRNA sequencing. Each case is presented with the proportion of correct or wrong predictions according to the number of votes obtained (2 or 3). The percentage is given by three votes confidence count only the result with three votes while the overall accuracy aggregates the 2 and 3 vote results. When combining results from both cases, the percentage of inconclusive results and the number of correct or wrong predictions are stated. Two different mapping parameters were tested, allowing respectively 20% of mismatches (part A) or 10% of mismatches (part B)

The predictions with the three votes using default mapping parameters (“relax mapping”) are very trustworthy as the accuracy is higher than 86% and 95% for cases 1 and 2, respectively. These promising results are obtained on the fraction of the elements representing 25-50% and 48-84% for case 1 and case 2, respectively. The remaining elements are classified with two votes (more information is available in Supplementary File 3). The prediction accuracy with two votes is much lower, whatever the case. So, knowing the number of votes obtained by each element is crucial when the results need to be interpreted (and this number is always given in the report generated by Cont-ID). For case 1, most elements were predicted with two votes meaning that one of the (three) rules had the opposite prediction, which might explain why the accuracy was lower. While for case 2, the majority of the elements were predicted with three votes. The explanation is probably in the “expected Infection/contamination” row in **Table 3A**: for all batches, there is more contamination than infection (from 28 infections for 77 contaminations – batch A to only 19 infections for 109 contaminations - batch D). As stated above in the text and **Figure 3**, Case 2 is designed to favour contamination detection at the expense of infections occurring at a low concentration that tend to be considered contamination (FN). Nevertheless, as a direct consequence, true contamination (TN) detection is high (see confusion_matrix in Supplementary File 3).

**Figure 3:**
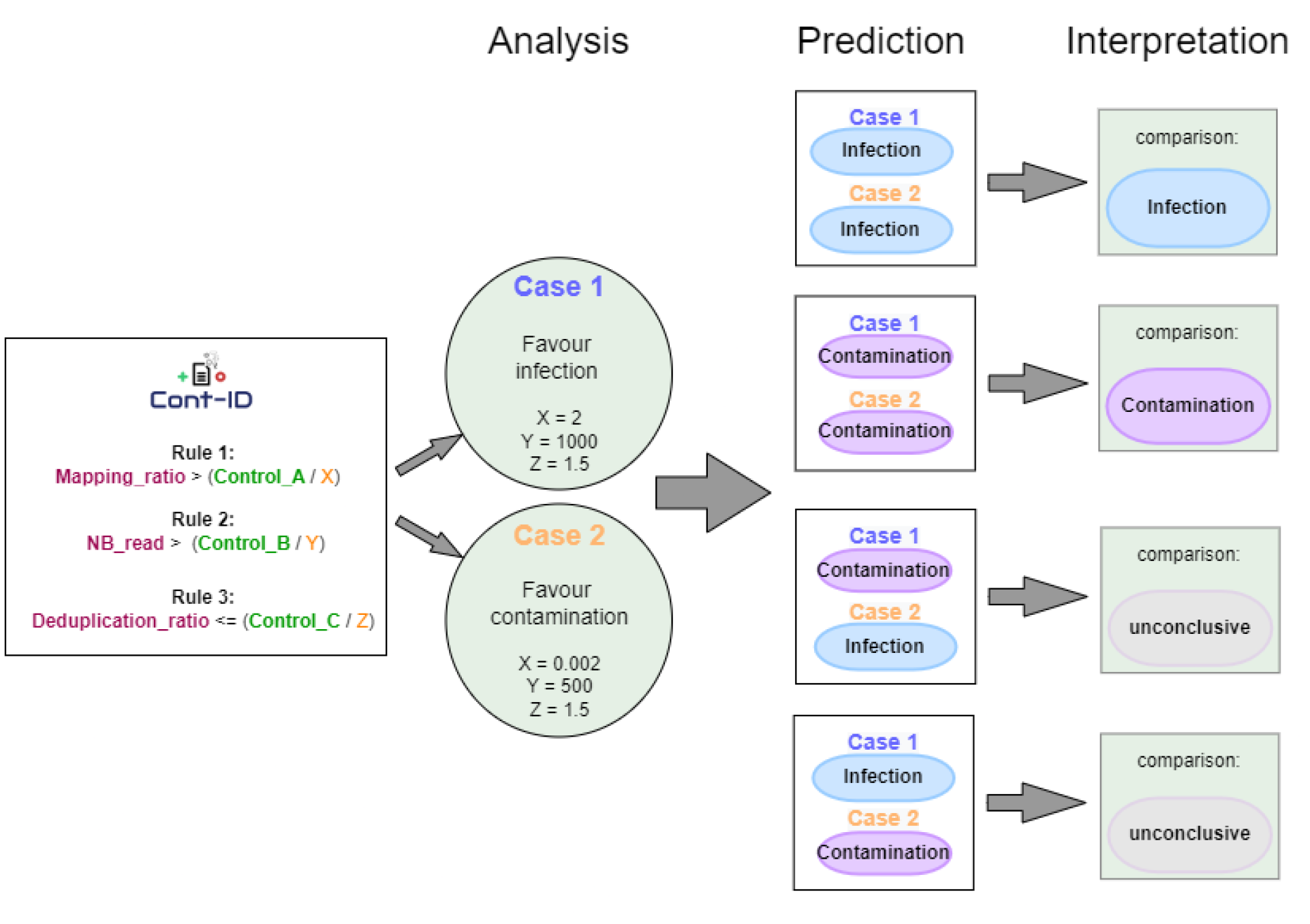
Cont-ID prediction when using the two default cases.

Overall, case 2 presented a higher accuracy (85-91% relax mapping) than case 1 (66-85%), while the combination of the two cases reached a similar one (82-95 % relax mapping). Those good results from combination accuracy mean that very few predictions are wrong (5 – 13) in both cases, but 16-33% of elements are not counted in the accuracy percentage because they are inconclusive. The combination’s importance relies on maintaining a high accuracy while highlighting the inconclusive prediction to prioritise them for manual expertise.

### The mapping parameters impacted the input files and the Cont-ID performance

In **Table 3**, we explored the impact on the prediction of two levels of mismatch tolerance (20% and 10%) when mapping the sequencing reads on the viral genome DB. The goal was to explore if changing a parameter from the primary bioinformatics step delivering the input files of Cont-ID could have an impact on the prediction. Strict mapping tends to lower the total number of elements tested due to a decrease in the number of samples for which we have very few reads mapped to a candidate virus. Cont-ID has more samples to process with a relaxed mapping, which should be better for threshold calculation. Logically, the elements lost by the strict mapping parameter should be predicted as “contamination” and present a relatively low number of reads. Indeed, those elements are most likely more distant reads (between 20 and 10 % mismatch with the reference genome) mapped on the virus. They could correspond to non-viral reads wrongly mapped in datasets from samples tested negative by classical indexing. For example, for batch B, on the 23 differential mapping results, the number of reads mapped ranged from 1 to 10. Of those 23 elements, 21 are classified as “contamination” the two remaining are labelled inconclusive. In batch C, there are 50 differential elements between the two parameters, with 37 correct (13 from BSV), 11 inconclusive (10 from BSV) and 2 wrong (2 from BSV) classified elements. In total, 25 elements (on the 50 - batch C) are from the non-integrated virus, of which 24 are labelled contamination (one inconclusive). The separation between integrated and non-integrated viruses is explained in **Table 4** and another publication (5).

**Table 4:** Percentage of the accuracy of case 1 and case 2 analysed alone or in combination on banana samples sequenced by ribodepleted totalRNA sequencing with relaxed mapping parameters. Each case is presented with the proportion of correct or wrong predictions according to the number of votes obtained (2 or 3). The percentage given by three votes confidence count only the result with three votes while the overall accuracy aggregates the 2 and 3 vote results. When combining results from both cases, the percentage of inconclusive results and the number of correct or wrong predictions are stated. Two types of viruses were tested, Non-integrated virus (part A) or integrated virus (part B).

The virus biology can impact Cont-ID performance: the case of virus integration in the host genome
| Batch | A- Non-integrated Virus |  |  |  | B- Integrated Virus (BSV) |  |  |  |
| --- | --- | --- | --- | --- | --- | --- | --- | --- |
|  | A | B | C | D | A | B | C | D |
| Sequencing method | TotalRNA |  |  |  | TotalRNA |  |  |  |

| Relax<br>mapping | Total element tested |  | 40 | 31 | 51 | 55 | 65 | 62 | 92 | 73 |
| --- | --- | --- | --- | --- | --- | --- | --- | --- | --- | --- |
|  | Expected<br>Infection/Contamination |  | 19/21 | 9/22 | 22/29 | 10/45 | 9/56 | 5/57 | 12/80 | 9/64 |
|  | Case 1 | 3 votes accuracy | 100% | 91% | 96% | 100% | 82% | 83% | 68% | 54% |
|  |  | 2 votes accuracy | 94% | 70% | 75% | 100% | 47% | 56% | 43% | 78% |
|  |  | overall accuracy | 98% | 77% | 94% | 100% | 58% | 61% | 50% | 74% |
|  | Case 2 | 3 votes accuracy | 100% | 85% | 100% | 100% | 97% | 100% | 100% | 93% |
|  |  | 2 votes accuracy | 50% | 91% | 82% | 67% | 62% | 58% | 89% | 67% |
|  |  | overall accuracy | 88% | 87% | 92% | 96% | 83% | 84% | 93% | 88% |
|  | Cases<br>combination | accuracy % | 100% | 88% | 96% | 100% | 79% | 79% | 92% | 91% |
|  |  | inconclusive % | 15% | 16% | 6% | 4% | 28% | 23% | 48% | 25% |
|  |  | correct<br>prediction | 34 | 23 | 46 | 53 | 37 | 38 | 44 | 50 |
|  |  | wrong prediction | 0 | 3 | 2 | 0 | 10 | 10 | 4 | 5 |
|  |  | inconclusive<br>(occurrence) | 6 | 5 | 3 | 2 | 18 | 14 | 44 | 18 |

Using the relaxed mapping parameter seems beneficial for prediction as the accuracy is better (82-95% relax, 78-90% strict). Moreover, thanks to the combination strategy, we can focus on the proportion of inconclusive; it is uneven with an important increase, 5% (strict) to 23% (relax) for batch A, while in batch D, it decreases from 28% to 16 %. However, when we look closely at the accuracy improvement, most comes from differential elements (present only with relax) that are ‘obvious’ contamination with few reads. So, most of the accuracy improvement did not come from very informative elements, except in some rare occurrences where it helped classify well elements in relax parameters that were inconclusive with strict or classified inconclusive elements in relax that were wrong with strict parameters. As an example, in batch C, on the 24 elements for BanMMV, BBRMV, BBTV and CMV common in both conditions (relax and strict), elements prediction is improved (from inconclusive [strict] to correct [relax]) for three of them (sample 3B1, 3B2 and 3B14 with BanMMV).

There is, therefore, a slight improvement with relaxed mapping parameters, and we set these parameters by default to generate the input files. Indeed, with the relaxed parameters, the number of reads for each element (including alien) increases along the rise of the number of elements in the batch. This means that we change the rule’s threshold (see Figure 2), which is critical for the threshold calculation in a way that seems more representative of reality than strict mapping. In these batches, some element metrics are very close to the threshold used for the rules and slightly changing those metrics or the alien metrics (the alien control metrics are obviously changed by the mapping parameters) can modify the prediction.

As we did not know the divergence of the virus genomes between different samples and the reference genomes, it seemed more logical to use relaxed mapping parameters by default. According to the virus system the user is working on and the ICTV demarcation criteria that go with it, these parameters should or could be adapted.

### The virus biology can impact Cont-ID performance: the case of virus integration in the host genome

To highlight the potential impact of the virus biology on the results of Cont-ID, the analysis of banana batches was split between integrated and non-integrated viruses. Indeed, several species of BSV are integrated into its host genome, which complicates the reliable detection of BSV infection from sequencing datasets. Consequently, it has been recommended to confirm any detection of BSV reads by an independent PCR test combined with immunocapture of viral particles (5).

Table 4 shows better accuracy and a lower proportion of inconclusive results for non-integrated viruses compared to BSV. More elements with contamination status are obtained when looking for BSV than non-integrated viral species. This over-representation of contaminants might be caused by the transcription of integrated sequences of BSV even without viral particles, which will raise the number of detected reads. These are two points that reduced the efficiency of our method on BSV and, by extension, might also concern any other viral species integrated into the host genome and able to produce transcripts.

The global accuracy is lower for BSV species (79-92%) compared to the other viruses (88-100%), even if the maximum accuracy obtained with batch C (92%) was high. In addition, the proportion of inconclusive results should also be considered, and this proportion was much higher for BSV (23-45%) than for the other viruses (4-16%). So, the overall performance of Cont-ID is lower when applied on BSV and did not solve the issues of appropriate detection in sequencing data of viral infection from viruses integrated into the plant genome. Consequently, BSGFV, BSIMV, BSMYV, and BSOLV, which correspond to different but closely related species of Banana streak virus (BSV) integrated into the *Musa* genome, were excluded from the calculation of performance criteria for the banana datasets. BSCAV was also excluded (despite not being integrated) because of its similarity with other BSV species.

### Performance of Cont-ID on diverse datasets

The performance of Cont-ID using the two cases was further evaluated while diversifying the hosts (fruit trees, grasses, humans) and the sequencing protocols (total RNA, small RNA, dsRNA).

**Table 5** shows the method’s accuracy on all datasets with relaxed mapping parameters (except for small RNA, see method). Overall, the accuracy of Cont-ID was 94%, with 15% of inconclusive results. The sRNA dataset provides a poor accuracy (50%) with 20% inconclusive; this can be explained by the (almost) absence of contamination (Expected Infection/Contamination 19/1) by the low level of reads found (see Supplementary Files 2 & 3 for more information). Apart from small RNA, the worst accuracy (88%) has been obtained from the batch B sequencing dataset of banana. Noteworthy, this protocol was independently evaluated for virus testing in banana, but its performance for virus detection was much lower than total RNA sequencing (5). The accuracy calculated from the single batch of dsRNA, with only 9 samples and 12 elements, was 100%. Even if not enough representative dataset was used for dsRNA, the method accuracy seems not too far from what we obtained in Total RNA, indicating that, Cont-ID is independent of the extraction method. On Total RNA, for banana samples, the accuracy ranged from 88% to 100%, with 4% to 15% of inconclusive results. The accuracy of the plant mix (G) was also very high (98%), with 8% of inconclusive results. On human datasets, the accuracy remained high (92-93%), but the inconclusive results reached up to 15 - 29%. Overall, the application of Cont-ID on human datasets reached similar performance in accuracy; the slightly worse inconclusive metrics can be explained by the fact that the adaptability metrics might not be the best ones for the human dataset and underlined the importance given at Cont-ID for the flexible adaptation of metrics and parameters.

**Table 5:**
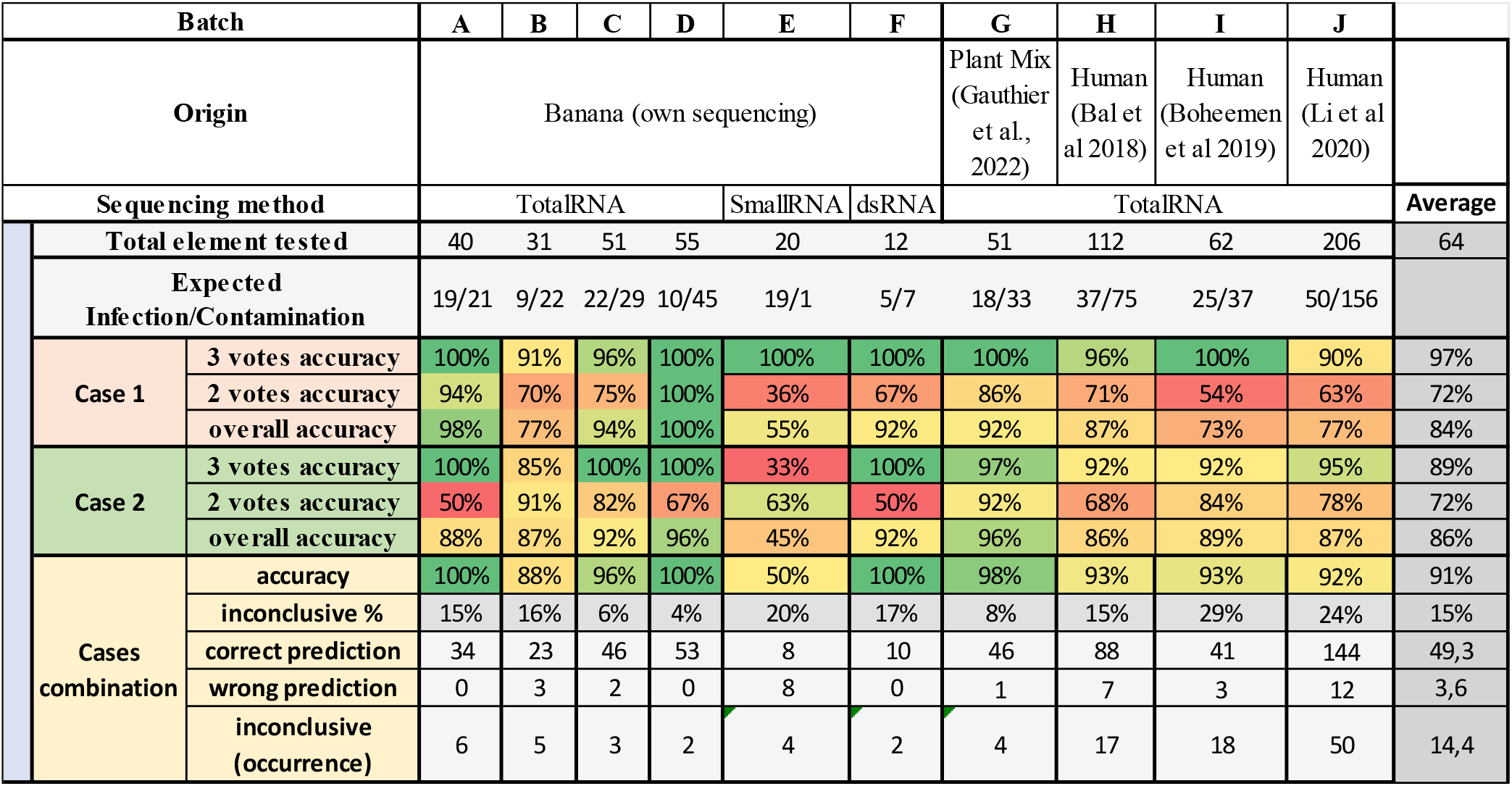
Percentage of the accuracy of case 1 and case 2 analysed alone or in combination from sequencing with relaxed mapping parameters (except for small RNA). Each case is presented with the proportion of correct or wrong predictions according to the number of votes obtained (2 or 3). The percentage given by three votes confidence count only the result with three votes while the overall accuracy aggregates the 2 and 3 vote results. When combining results from both cases, the percentage of inconclusive results and the number of correct or wrong predictions are stated. Several virus datasets were tested, banana samples (only non-BSV viruses are considered), a mix of plants, and human datasets.

For most of the datasets, case 1 performed worse than case 2, probably due to the design of the case metrics (see **Figure 2**), where case 1 values were determined to favour infection. The expected infection/contamination ratio showed that for all the datasets but E (small RNA), there was a lot more contamination than infection; therefore, case 1 overpredicted infection, lowering its accuracy. In the E dataset, case 1 (55%) performed better than case 2 (45%) as expected; it is also the case for the human dataset H (97% case 1, 96% case 2), even if the ratio (37/75) leans toward contamination.

Those results indicated that Cont-ID performed well in classifying cross-contamination in very different virus-host systems, even if some adjustments may be needed in some cases in the future. The different levels of flexibility of Cont-ID made such adjustments possible. To provide an example of analysis, all the information regarding batch C from the input file to the analysis file (including raw results) is available in Supplementary File 3.

## Discussion

Despite significant efforts to limit cross-contamination (dual indexes, inter-run washing …), this still represent a concern and the appropriate distinction between low-level infection, and cross-sample contamination is crucial for the large-scale development of HTS technologies as a diagnostic test. Furthermore, it should be adequately managed because identifying and monitoring the cross-contaminations improves the detection results’ reliability. In other words, it can help to find the source of contamination in the laboratory, take appropriate measures to minimise it, and raise confidence in the detected viruses.

This publication improved a preliminary work on determining an adaptative contamination threshold for the detection of plant viruses (5), which uses the maximal number of alien virus reads contaminating a sample as the threshold of detection for each sequencing batch. So, instead of using a fixed number for the contamination threshold as done in the literature, the threshold is adapted to the level of contamination monitored in the batch thanks to the alien control. The former publication used a single threshold corresponding to the maximum number of alien virus reads in a sample. Some limitations of this previous threshold, for example, overestimating contamination when viral reads are in low number for a virus, underlined the need for improvements. This was achieved with Cont-ID through the definition of multiple formal rules, the automation of calculation and the ability to adapt the thresholds and rules by the user. The tool’s prediction relies on basic and usual information generated by bioinformatics analysis of sequencing data (mapping and duplication numbers) and the use of external alien control. The criteria based on reads (relative) abundance of each virus in each sample and the (approximation of) number of identical reads for a virus between samples performed well while being relatively easy to generate. Our objective with this tool was to show that exploring data generated by standard bioinformatic procedures can facilitate the identification of cross-contamination between samples.

Cont-ID discriminated virus infection and cross-contamination between samples with a global accuracy of 91 % (median=95%) on the diverse range of datasets included in its evaluation. The diversity of situations included viral species belonging to diverse viral families with cellular hosts belonging to plant or animal kingdoms and three different library preparation protocols. Importantly, the default values of adaptability metrics determined from banana dataset predicted cross-contamination with high accuracy (96%, on banana excluding small RNA) and remain high even on human datasets (94%). To further help the user in the analysis, we provide the detailed votes prediction in the result file (see Supplementary File 3). This is, therefore, a solid basis for the diagnostician to check the level of confidence in the generated results. Indeed, each prediction made by the method uses at least two rules to determine the classification of the element for each case. A prediction with three votes is more confident than with two votes. But all predictions with two votes do not provide the same confidence as it depends on which rules predicted what. Of our three rules, two rely more or less directly on abundance estimation, which means that when that metric is not obtainable in a reliable way, the tools’ predictions will be impacted, and predictions with those rules might be less confident. On the other hand, rule three (deduplication ratio) is less effective when the read numbers are low. Depending on the scenario, the user should consider the relative confidence of each rule when trying to confirm Cont-ID prediction. This underlines again the importance of proper interpretation of the obtained results based on the virus biology

The prediction quality depends on the input data quality, meaning that the deduplication and mapping parameters are essential and should be carefully considered while evaluating their impact on the results. For example, some deduplication tools remove reads if a (small) read is contained in another (larger) read; having that option active or not will significantly impact the deduplication ratio. As shown in the results, mismatch parameters are very impactful for the mapping. Considerations like ICTV demarcation criteria or what parameters the biologist would use to reconstruct the whole viral genome are helpful in deciding the ones to use for Cont-ID input. In that regard, testing and expertise in bioinformatics analysis are heavily beneficial. Here, the 20% mismatch parameter performed well; it might be different in other datasets (viral composition) configurations or when working with databases containing many reference genomes. Indeed, independently of the mismatch parameters used, using more genome references for each expected species could also improve the ability to detect sequences from distant isolates by better covering the genetic diversity of the virus.

The biology of the virus should also be considered, as shown by the results obtained with viruses with functionally integrated genomes in the host, like BSV species. Our conclusion is that they should be considered independently from the non-integrated viruses. It was challenging to extract a reliable metric for BSV as the differentiation between reads from integrated genomes and reads from viral particles is impossible. Indeed, the biology of viruses integrated into the host genome differs from non-integrated viruses, as viral genome transcription can happen without viral particle production. We have not tested our method on species with different biological behaviour like viroids or phages. But optimisation of the adaptative metrics might likely be required in order the use Cont-ID with high accuracy. Viroid genomes are generally smaller than viruses, while the phage genome tends to be much larger and has specific biological features. For example, a different level of identical reads and abundance (calculation based on reads number) could be obtained between the different scales of genome size.

For these reasons, Cont-ID allows the evaluation of other values for adaptability metrics (X, Y, Z) by each user to adapt the tool and optimise its diagnostic performance depending on the biological matrix, the protocol and the purpose of the test. Independently, the user can also adapt the metrics to reach the appropriate balance between FN and FP by deciding if, for the purpose of the test and the available resource for confirming detection, it is preferable to be overpredicting contamination to be confident that all the virus detection remaining are true infection or the opposite (overpredicting infection to be sure not to miss any).

In our tests, the analysis of the wrong predictions showed that none of the proposed rules (and adaptability metrics values) allowed us to reach satisfactory accuracy with a proper balance between FN and FP (see Supplementary File 1). We have observed that using two sets of adaptability metrics (one to favour contamination and the other, infection prediction) gave a higher accuracy. In a real scenario (with infection status not known for the samples), it is difficult to know if HTS virus detection (at low concentration) is in the majority due to true infection or cross-contamination. The two-case strategy allows the biologist to predict both scenarios with at least one case accurately. Indeed, if the expected ratio of infection/contamination is unknown, the relative performance of cases 1 or 2 will be unknown, so it seems preferable to use the combination results instead of the individual.

If both cases agree, the assumption is that the prediction is correct. Nevertheless, combining the results will provide a list of interesting inconclusive results. Each inconclusive result means that the two cases delivered opposite predictions. Therefore, the scientist should address those results when analysing Cont-ID prediction by checking the number of rules for each prediction, for example, knowing if 2 or 3 rules agreed and checking the results of each rule: How close to the threshold was the read abundance/ratio and/or the duplication rate? Spotting the few errors that may occur requires excellent manual expertise as the usual manual verification methods may also indicate the wrong decision (if there are many reads from cross-contamination, the mapping results can be wrongly positive while the and/or the (RT)-PCR can also be wrongly positive if the contamination occurred at an early stage and the (RT)- PCR was carried out on the same nucleic acids extract). Other information about the virus-plant interaction should be considered, like virus-species-cultivar compatibility or geographical virus distribution (see investigation on unexpected viruses (5)).

Cont-ID also presents some limitations that need to be discussed. First, the number of identical reads estimation comes from the deduplication procedure, which is an approximation, and that can be a problem because it can consider the non-specific reads (reads that are not coming from cross-contamination but that are identical to another sample from a common area of the genome) as identical to the probable source of contamination by mistake. Indeed, this can be the case if, for example, two samples are infected by the same virus isolate at very different concentrations. The presence of duplicated reads might suggest contamination instead of a low-level infection. The risk of such an extreme situation is limited using two other rules, although interpreting the data will require good expertise in virus genomic variability and detailed information on the sample origins and virus prevalence and diversity.

In addition, the duplication metric assumes that contamination (if any) comes from the sample with the highest number of reads. This theory seems logical since the more reads in a sample, the higher the probability of detecting a few reads from it contaminating other samples (potential of contamination). Nevertheless, it can create a bias when a virus is highly abundant in two (or more) samples and detected with a low frequency in others. In that case, it is difficult to determine the true origin of cross-contamination. Such a case could be a fundamental limit of our current method. If several samples with a very high abundance of reads are present in a batch, as developed here, Cont-ID should be applied as many times as the number of highly abundant samples. Ideally, Cont-ID should include the read duplication comparison of each sample to all other samples for a virus, but this can raise additional issues (like contamination from several origins at the same time), and, at this stage, it was not implemented.

We must also keep in mind that the relative quantity of genetic material between samples might change because the biologist normalises the quantity of DNA/RNA at two steps of the process: before starting library preparation and during the pooling of the prepared libraries. Meaning that the differential in genomic material concentration (potential of contamination of a sample) is resettled. If cross-contamination happens before that step, it can cause less accurate predictions from Cont-ID. This bias in the estimation of abundance is another limitation of our method.

Using an (alien) control helps to know the expected level of contamination but is also impacted by the limit of detection inherent to the standard bioinformatic procedures. Indeed, working with very few reads for some viruses makes some analyses impossible when below their detection limit. For example, the calculation of the duplication rate below a minimal number of reads (in this study, we chose 5) of a virus did not make sense. The limit of calculation of the input metrics is another limitation of Cont-ID.

Cont-ID accuracy was high, but additional improvements can probably be explored. For example, by exploiting the ability of other metrics generated during bio-informatic analyses (like RKPM, genome coverage percentage, relative coverage depth repartition, …) to help detect contamination. In fact, some of these metrics with several thresholds were tested for Cont-ID before selecting the three rules described in **Figure 2** that provided the highest accuracy (in both contamination and infection determination). Importantly, values leading to a perfect scenario were not identified, and a two-cases classification system was set up (more information in Supplementary File 1).

Nevertheless, adding more metrics will also complexify the decision system. If more metrics are considered for cross-contamination prediction, other implementations (decision tree, machine learning …) might be envisioned to replace the current voting system. On the other hand, the detection in the alien control of sequencing reads of other viruses detected in the tested samples is also the consequence of contamination from one of the tested samples toward the alien control. This information is not used now but could also be considered for future improvements as it requires less complex to implement. In addition, it might allow refinement of Cont-ID, potentially introducing an adaptation of threshold per virus instead of a single threshold for all samples from the sequencing batch. The idea is that two viruses present in the same batch may have different relative abundance behaviour in the samples, so setting up a limit that can adapt for each virus should improve the tool’s ability to distinguish real infection from cross-contamination. Finally, working with the combination of all/some viruses profile instead of each individually for contamination check (similarly to what is used in metabarcoding of bacteria) can also be considered. Indeed, when a sample contaminates another, it is expected that all the viruses (highly frequent) from the contaminating sample can be found in the contaminated samples. Monitoring the virus detection profile of samples can provide additional information for cross-contamination (and ease the quest for contamination origin). Even if there is still improvement to be made, Cont-ID has already delivered an excellent ability to consider the level of contamination genuinely present in the batch.

In conclusion, detection of cross-contamination is complex; in the age of sequencing, the contaminant issue is increasingly important; therefore, Cont-ID will facilitate the interpretation of results by the virologist/diagnostician and reduces the confirmation burden. We demonstrated that simple metrics like relative abundance estimation and redundancies of genetic material (reads duplicates) could help monitor contamination occurring in the laboratory. The method accurately distinguished cross-contamination from infection in very diverse HTS viral datasets. Our standard parameters allowed very good accuracy (median = 95%); in addition, Cont-ID has several levels of flexibility and can be adapted by each user to take into account the specificities of the detection test (purpose of the test, type of samples, viruses to be detected, laboratory work, available resources…). We believe this is the first significant step toward increasing the monitoring and management of sample cross-contamination when using HTS technologies for virus detection.

## Availability and requirements

Project name: Cont-ID

Project home page: https://github.com/johrollin/Cont_ID

Operating system(s): Platform independent

Programming language: Python (v3.7)

Other requirements: pandas; NumPy

License: GNU GPL-3.0

Any restrictions to use by non-academics: none

## Declarations

### Ethics approval and consent to participate

Not applicable.

### Consent for publication

Not applicable.

### Availability of data and materials

The datasets generated and/or analysed during the current study are available in the NCBI Sequence Read Archive (SRA) repository at https://www.ncbi.nlm.nih.gov/sra (See Table 1). In addition, cont-ID is freely available and can be downloaded with the following command (without <>): <https://github.com/johrollin/Cont_ID>. It can be used as a command line application on a personal computer on any operating system (Linux, MacOSX or Windows) with python.

### Authors’ contributions

WR and SM designed the data preparation and sequencing procedure. WR, JR and SM designed the bioinformatics analysis. JR implemented the program. JR ran CroCo analyses. WR and JR validated Cont-ID results with previous PCR indexing results and re-analysed outlier. JR and SM drafted the manuscript. All authors read and approved the final manuscript.

## Acknowledgements

We thank Angelo Locicero for technical support and Gladys Rufflard for administrative support. Delphine Masse (ANSES, La Réunion, France), Kathy Crew and John Thomas (Queensland Alliance for Agriculture and Food Innovation, Brisbane, Australia), Mathieu Chabannes and Marilyne Caruana (CIRAD, Montpellier, France) are also acknowledged for kindly providing Musa reference samples. Special thanks to Marie-Emilie Gauthier and Roberto Barrero for providing dataset G and discussing cross-contamination in viral metagenomes with us.

